# Self-activation of Wnt signaling in pre-granulosa cells is required for ovarian folliculogenesis

**DOI:** 10.1101/2020.10.20.346304

**Authors:** Okiko Habara, Catriona Y. Logan, Masami Kanai-Azuma, Roeland Nusse, Hinako M. Takase

## Abstract

In mammalian ovaries, immature oocytes are reserved in primordial follicles. Precise control of primordial follicle activation (PFA) is a prerequisite for proper reproduction. Although Wnt signaling is thought to be involved in folliculogenesis, the timing and function of Wnt activity remain unclear. Here we show that canonical Wnt signaling is pivotal for the differentiation of pre-granulosa cells (pre-GCs) and subsequent oocyte maturation during PFA. We identified several *Wnt* ligands expressed in pre-GCs that cell-autonomously function via canonical Wnt activity. Inhibition of Wnt ligand secretion from pre-GCs/GCs led to infertility due to impaired pre-GC differentiation, whereas constitutive stabilization of β-catenin induced thickening of the pre-GCs. Our data support a two-step model of PFA in which self-activation of Wnt signaling promotes the transition of pre-GCs to GCs, and mature GCs then support oocyte reawakening. We anticipate that application of Wnt inhibitors or activators in vitro will lead to improved fertility treatments.

## Introduction

In female mammals, including humans, precise control of folliculogenesis is essential for fertility. Oocytes are protected and grow within follicles, which are the fundamental units of the ovary. Dormant oocytes are arrested at the diplotene stage of Meiosis I, reserved in primordial follicles, and surrounded by pre-granulosa cells (pre-GCs) (Pepling, 2006; Pepling and Spradling, 2001). Only a small proportion of primordial follicles is activated concurrently, with activation resulting in follicle growth and the serial development of primary, secondary, and antral follicles. Although primordial follicles are able to survive for years to decades, once activated their lifespan is limited to days to months, with their potential fates being either ovulation or atresia (McGee and Hsueh, 2000). In women with primary ovarian insufficiency (POI), the number of follicles declines rapidly and menopause occurs before age 40, resulting in severe fertility problems. POI is not a rare condition, with an estimated prevalence of 1% of women. The cause of POI remains unknown in most cases, but misregulation of primordial follicle activation (PFA) is thought to be a contributing factor (Jankowska, 2017; De Vos et al., 2010). Precise control of PFA is thus required for maintenance of female reproductive ability, with new oocytes not being thought to be generated after birth (Lei and Spradling, 2013).

Given that functional gonadotropin receptors are not present in primordial follicles, PFA is thought to be controlled in a gonadotropin-independent manner (Mason et al., 1986). PFA is characterized morphologically by oocyte growth to a diameter of >20 μm and conversion of the squamous pre-GCs into cuboidal GCs. Two key events thus proceed synchronously during PFA: oocyte outgrowth and the proliferation-differentiation of pre-GCs (Adhikari and Liu, 2009). Several intracellular signaling pathways in oocytes have been implicated in control of their dormancy or growth. The transcription factor FoxO3 (Forkhead box O3) and the phosphatase PTEN (phosphatase and tensin homolog deleted from chromosome 10) are required for the quiescence of oocytes, whereas activation of the phosphoinositide 3-kinase (PI3K)–Akt– mammalian target of rapamycin (mTOR) pathway contributes to oocyte reawakening (Adhikari et al., 2009; Castrillon et al., 2003; John et al., 2008; Reddy et al., 2008). In addition, environmental factors such as hypoxia and mechanical stress can influence maintenance of the dormant state of oocytes (Nagamatsu et al., 2019; Shimamoto et al., 2019). In contrast, the mechanism underlying the activation of pre-GCs has been less well understood. Differentiation of pre-GCs was found to be disrupted in mice deficient in FoxL2 or both GATA binding protein 4 (GATA4) and GATA6, although such gene ablation from the early embryonic period might also affect GC lineage identity (Padua et al., 2014; Schmidt et al., 2004). Activation of pre-GCs is likely to trigger oocyte reawakening, given that the associated activation of the mTOR pathway in pre-GCs results in the production of Kit ligand (KitL), which contributes to oocyte activation (Liu et al., 2014). However, a comprehensive understanding of the mechanism of PFA requires clarification of how GCs differentiate during this process.

Wnt signaling is an evolutionarily conserved system for cell-cell communication that can be classified broadly into canonical (β-catenin–dependent, also referred to as Wnt/β-catenin signaling) and noncanonical (β-catenin–independent) pathways (Martin-Orozco et al., 2019). Nineteen Wnt ligands have been identified and contribute to diverse processes such as development, stem cell control, and disease in mice and humans (Nusse and Clevers, 2017; Wiese et al., 2018). Whereas Wnt4-mediated canonical Wnt signaling has been shown to be important for sex determination during embryonic development (Parma et al., 2006; Vainio et al., 1999), the function of Wnt signaling during postnatal folliculogenesis has remained unclear. Both Wnt2 knockout and GC-specific Wnt4 knockout female mice were found to manifest a slight reduction in fertility, with the mild nature of this defect in each case likely being due to functional redundancy among Wnt ligands (Boyer et al., 2010; Monkley et al., 1996). More recently, oocyte-derived R-spondin 2 (Rspo2) was shown to contribute to the activation of Wnt signaling in GCs (De Cian et al., 2020). Rspo2 is a Wnt agonist that is secreted extracellularly and enhances canonical Wnt signaling (Kazanskaya et al., 2004), and follicle growth was found to be impaired in ovaries with loss of Rspo2 function by transplant experiments(De Cian et al., 2020). Although Wnt signaling is implicated together with other important factors such as GDF9 (growth differentiation factor 9) and BMP15 (bone morphogenetic protein 15) in PFA (Dong et al., 1996; Dube et al., 1998), its mechanism of action has been unknown. Here we reveal an essential role of canonical Wnt signaling in regulation of the transition of pre-GCs to mature GCs during PFA by focusing on postnatal folliculogenesis and taking advantage of mouse mutants that avoid the issue of the redundancy of Wnt ligands.

## Results

### Canonical Wnt signaling in pre-GCs is essential for female fertility

Although Wnt signaling has been implicated in adult folliculogenesis, the spatiotemporal patterns of Wnt ligand expression in the mouse ovary have been insufficiently characterized (Harwood et al., 2008). Both the complexity and specificity of Wnt signaling in mice are due in part to the expression of 19 Wnt ligands. To determine which Wnt ligands are expressed during folliculogenesis, we performed in situ hybridization analysis with ovaries from 3-week-old wild-type (WT) mice for all 19 *Wnt* mRNAs (Figure S1). Among the 19 Wnt ligands, the mRNAs for Wnt4, Wnt6, and Wnt11 were detected in the GC lineage from the primordial follicle to primary follicle stages (Figure 1A). The abundance of these Wnt ligand mRNAs gradually declined in association with the transition to secondary follicles. We did not observe intense expression of Wnt ligands in oocytes. These data thus suggested that pre-GC/GC–derived Wnt signals might contribute to the early stages of folliculogenesis.

**Figure 1.**
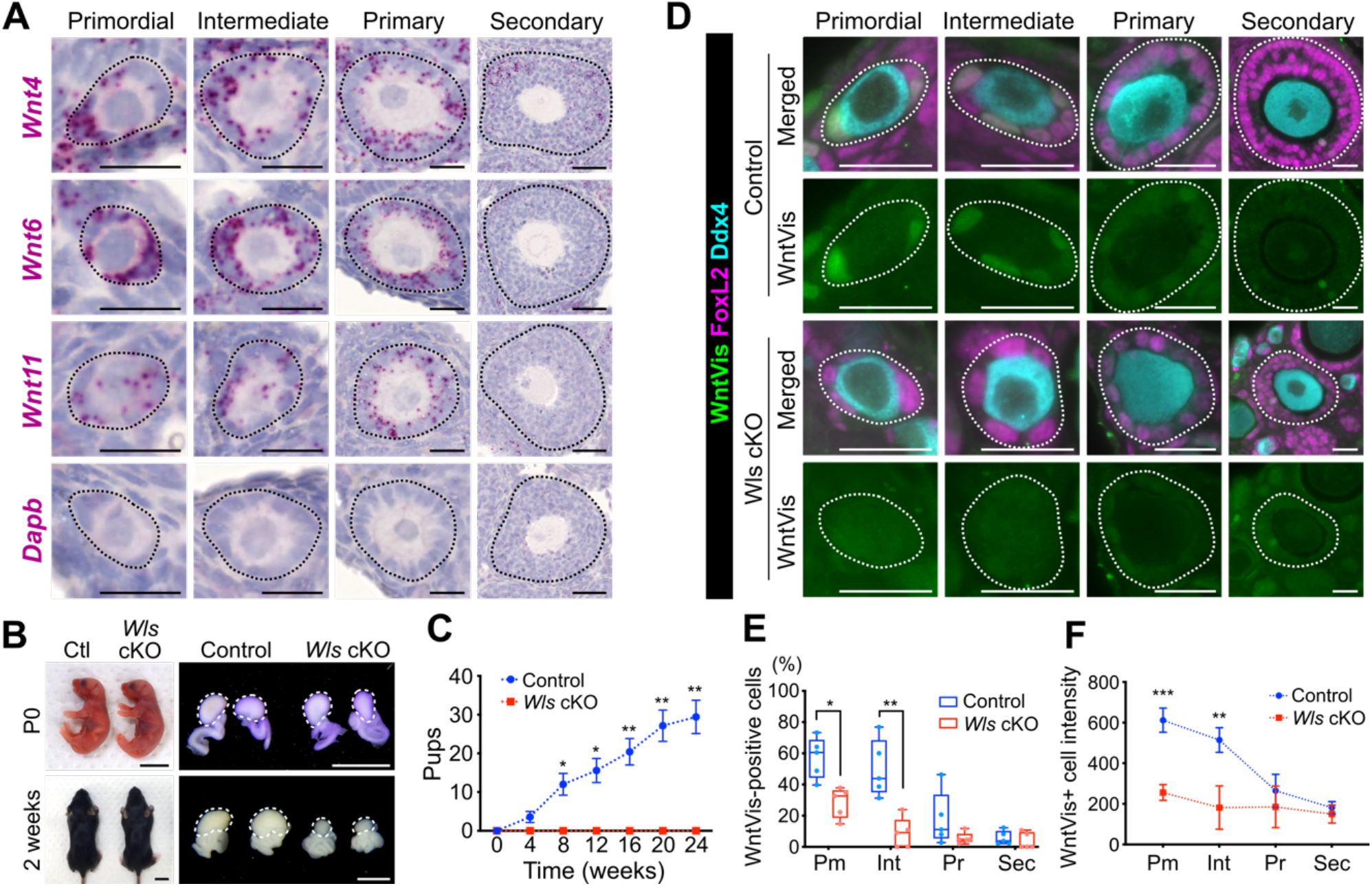
Canonical Wnt signaling in primordial follicles. (A) In situ hybridization analysis of *Wnt4*, *Wnt6*, *Wnt11*, and *Dapb* (negative control) mRNAs (red) in ovaries of 3-week-old WT mice. Follicles were classified as primordial, intermediate, primary, or secondary, and they are demarcated with dotted lines. Scale bars, 50 μm (rightmost panels) or 20 μm (other panels). (B) Gross morphology of the body and ovaries (indicated by dotted lines) of *Wls* cKO and littermate control (Ctl) mice at P0 and 2 weeks of age. Black scale bars, 1 cm; white scale bars, 100 μm. (C) Cumulative number of pups born to control or *Wls* cKO female mice housed with WT males for 24 weeks beginning at 8 weeks of age. Data are means ± SEM (*n* = 7 females of each genotype). \**P* < 0.05, \*\**P* < 0.01 (two-way ANOVA with Sidak’s test for multiple comparisons). (D) Immunofluorescence staining of WntVis (green), FoxL2 (magenta), and Ddx4 (cyan) in the ovaries of 3-week-old *Wls* cKO and littermate control mice harboring the *R26-WntVis* allele. The white dotted lines indicate follicles. Scale bars, 20 μm. (E) Percentage of WntVis-positive cells among pre-GCs/GCs for each follicle type (Pm, primordial; Int, intermediate; Pr, primary; Sec, secondary) as determined from images as in d. The boxes indicate the median and 25th and 75th percentiles, the whiskers indicate minimum and maximum values, and five mice of each genotype were examined. \*\**P* < 0.01, \*\*\**P* < 0.001 (unpaired multiple *t* tests with Holm-Sidak correction). (F) Fluorescence intensity of WntVis-positive cells for each follicle type. Data are means ± 95% confidence interval (*n* = 18 to 256 cells from five mice of each genotype). \*\**P* < 0.01, \*\*\**P* < 0.001 (unpaired multiple *t* tests with Holm-Sidak correction).

To examine the effects of attenuation of Wnt signaling, we generated ovarian somatic cell– specific *Wntless* conditional knockout (*Wls* cKO) mice by crossing *Sf1-Cre* mice (which express Cre recombinase under the control of the steroidogenic factor 1 gene promoter) to mice harboring a “floxed” (*Wls*^flox^) or ubiquitous deletion (*Wls*^del^) allele of *Wls*. Given that Wls is required for secretion of all Wnt ligands, the resulting *Sf1-Cre;Wls*^flox/del^ (*Wls* cKO) mice allow us to examine the effects of inhibiting Wnt ligand secretion specifically from ovarian somatic cells, including GC lineage cells, from embryonic day (E) 11.5 (Dhillon et al., 2006; Piprek et al., 2019). *Wls* cKO mice were born with a normal Mendelian frequency, and no obvious morphological abnormalities were apparent during development through adulthood (Figure 1B). The ovaries of *Wls* cKO mice were similar to those of littermate control mice at postnatal day (P) 0, whereas they manifested atrophy at 2 weeks of age (Figure 1B). To evaluate reproductive performance, we housed 8-week-old control or *Wls* cKO female mice (*n* = 7 per genotype) with WT males for 24 weeks. *Wls* cKO females were completely infertile (Figure 1C), even though they engaged in spontaneous mating behavior.

To identify Wnt-responding cells, we evaluated the ovaries of a Wnt signal reporter mouse line, *R26-WntVis*. The green fluorescent protein (GFP) reporter activity of these mice reflects the activity of the canonical Wnt signaling pathway (Takemoto et al., 2016). GFP was specifically expressed in the GC lineage from the primordial to primary follicle stages (Figure 1D), consistent with the expression pattern of Wnt mRNAs (Figure 1A). FoxL2 was examined as a marker for pre-GCs/GCs and Ddx4 as a marker for oocytes in this analysis. The WntVis signal was also sparsely detected in the interstitial cells, Theca cells, and ovarian epithelium, but not in blood vessels (Figure S2A–S2E). It was undetectable in oocytes (Figure 1D). The WntVis signals were most abundant and intense in pre-GCs of primordial follicles, and they became less abundant and less intense with follicle growth (Figure 1D–1F). Both the number of WntVis-positive pre-GCs/GCs and WntVis fluorescence intensity were significantly reduced in *Wls* cKO mice harboring the *R26-WntVis* allele compared with control mice (Figure 1D–1F). These results thus suggested that autocrine or cell-autonomous Wnt signaling activity in pre-GCs is required for female fertility.

### Wnt signaling is required for pre-GC to GC differentiation-maturation

To examine the cause of the ovarian defects of *Wls* cKO mice, we performed a more detailed morphological analysis (Figure 2A). Immunostaining of Ddx4 revealed that the number of oocytes per ovary did not differ significantly between *Wls* cKO and control mice at P0 (Figure 2C). Periodic acid– Schiff staining with hematoxylin (PAS-H) revealed few atypical follicles, such as those containing multiple oocytes, in the ovaries of the mutant females at 2 weeks of age (Figure 2B). Abnormal sexual differentiation was also not apparent, as confirmed by sex genotyping. These data suggested that germ cell survival during embryonic development, cyst breakdown, and sex determination were not affected in *Wls* cKO mice. Whereas differentiated cuboidal GCs were apparent in growing follicles containing oocytes with a diameter of 20 to 40 μm of control mice, flattened and morphologically abnormal GCs were detected in *Wls* cKO mice (Figure 2B). In contrast, no morphological abnormalities were detected in primordial follicles with an oocyte size of <20 μm (Figure 2B). Quantitative analysis revealed that the GC layer was significantly thinner in growing follicles of *Wls* cKO mice, whereas it was similar in primordial follicles of both genotypes (Figure 2D and 2E). GCs in *Wls* cKO mice were less likely to become multilayered, and even if they did form multiple layers the layers were uneven (Figure 2B, see Figure 3A). These results thus indicated that PFA-associated pre-GC differentiation is suppressed in the absence of Wnt signaling.

**Figure 2.**
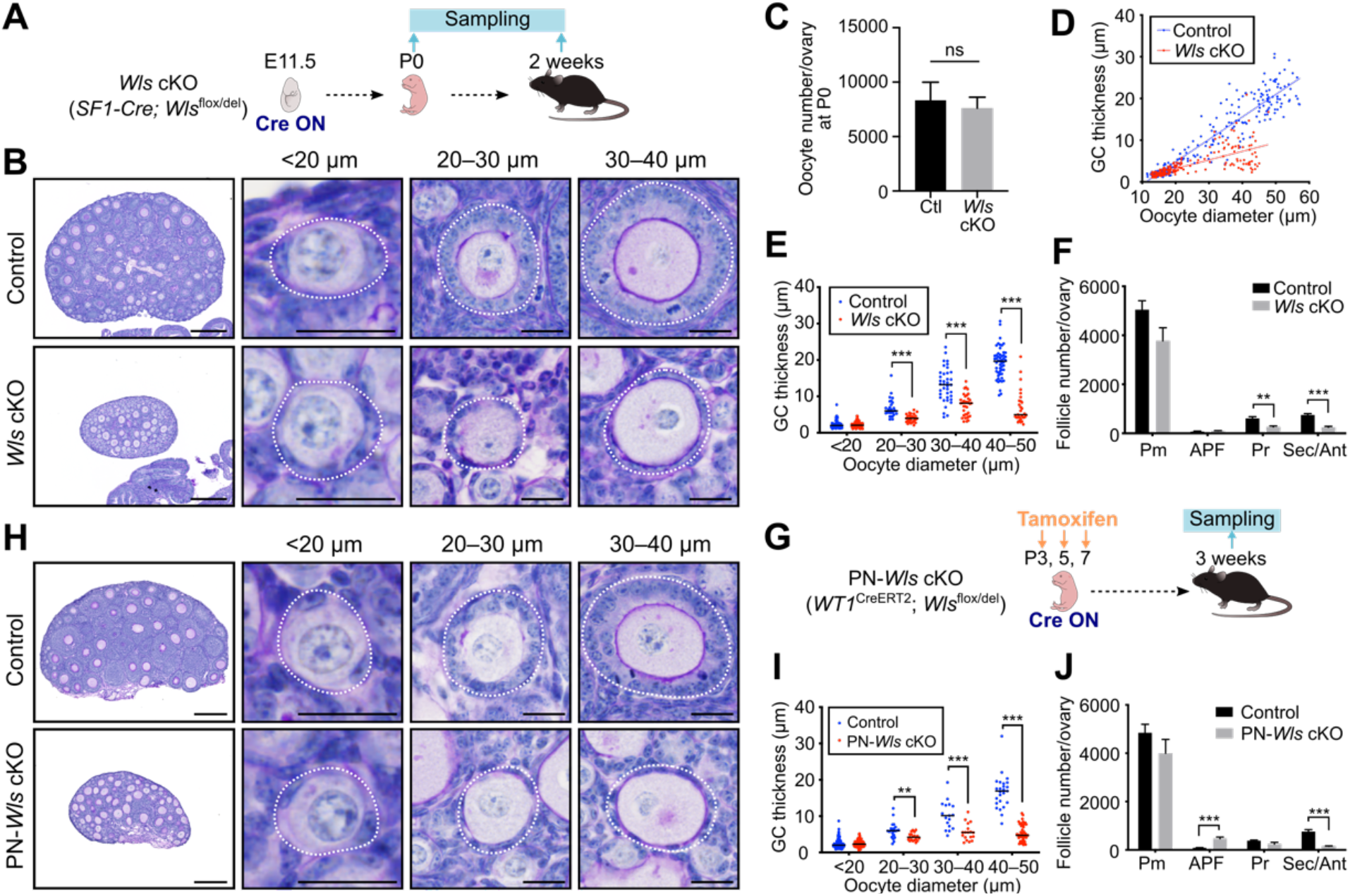
*Wls* cKO mice show an impaired transition of pre-GCs to GCs. (A and G) Experimental schemes for examination of *Wls* cKO (A) or PN-*Wls* cKO (G) mice to determine the effects of embryonic or postnatal deletion of *Wls* in ovarian somatic cells on folliculogenesis. (B and H) PAS-H staining of ovarian sections from 2-week-old *Wls* cKO mice (B) or 3-week-old (tamoxifen-treated) PN-*Wls* cKO (H) mice. Follicles were classified by oocyte size and are demarcated by white dotted lines. Scale bars, 200 μm (leftmost panels) or 20 μm (other panels). (C) The number of Ddx4^+^ oocytes per ovary of *Wls* cKO or littermate control (Ctl) mice as determined by immunohistochemical staining at P0. Data are means + SEM (*n* = 7 mice of each genotype). ns, not significant (nonparametric Mann-Whitney matched-pairs test). (D) Scatter plot for the distribution of oocyte diameter and GC layer thickness for follicles of *Wls* cKO (*n* = 436 follicles) and control (*n* = 412 follicles) mice at 2 weeks of age as determined from images similar to those in (B). Regression lines are included. (E and I) GC layer thickness categorized by oocyte diameter for follicles of *Wls* cKO mice at 2 weeks of age (*n* = 26 to 110 follicles from four mice of each genotype) (E) or tamoxifen-treated PN-*Wls* cKO mice at 3 weeks of age (*n* = 17 to 228 follicles from six mice of each genotype) (I). Horizontal lines represent the median. \*\**P* < 0.01, \*\*\**P* < 0.001 (unpaired multiple *t* tests with Holm-Sidak correction). (F and J) Quantification of follicle number per ovary for 2-week-old *Wls* cKO mice (F) or 3-week-old (tamoxifen-treated) PN-*Wls* cKO mice (J) as determined by immunohistochemical staining for Ddx4. Follicles were classified as primordial (Pm), activated primordial (APF: oocyte diameter of >20 μm without cuboidal GCs), primary (Pr), or secondary/antral (Sec/Ant). Data are means + SEM (*n* = 7 mice per genotype). \*\**P* < 0.01, \*\*\**P* < 0.001 (unpaired multiple *t* tests with Holm-Sidak correction).

**Figure 3.**
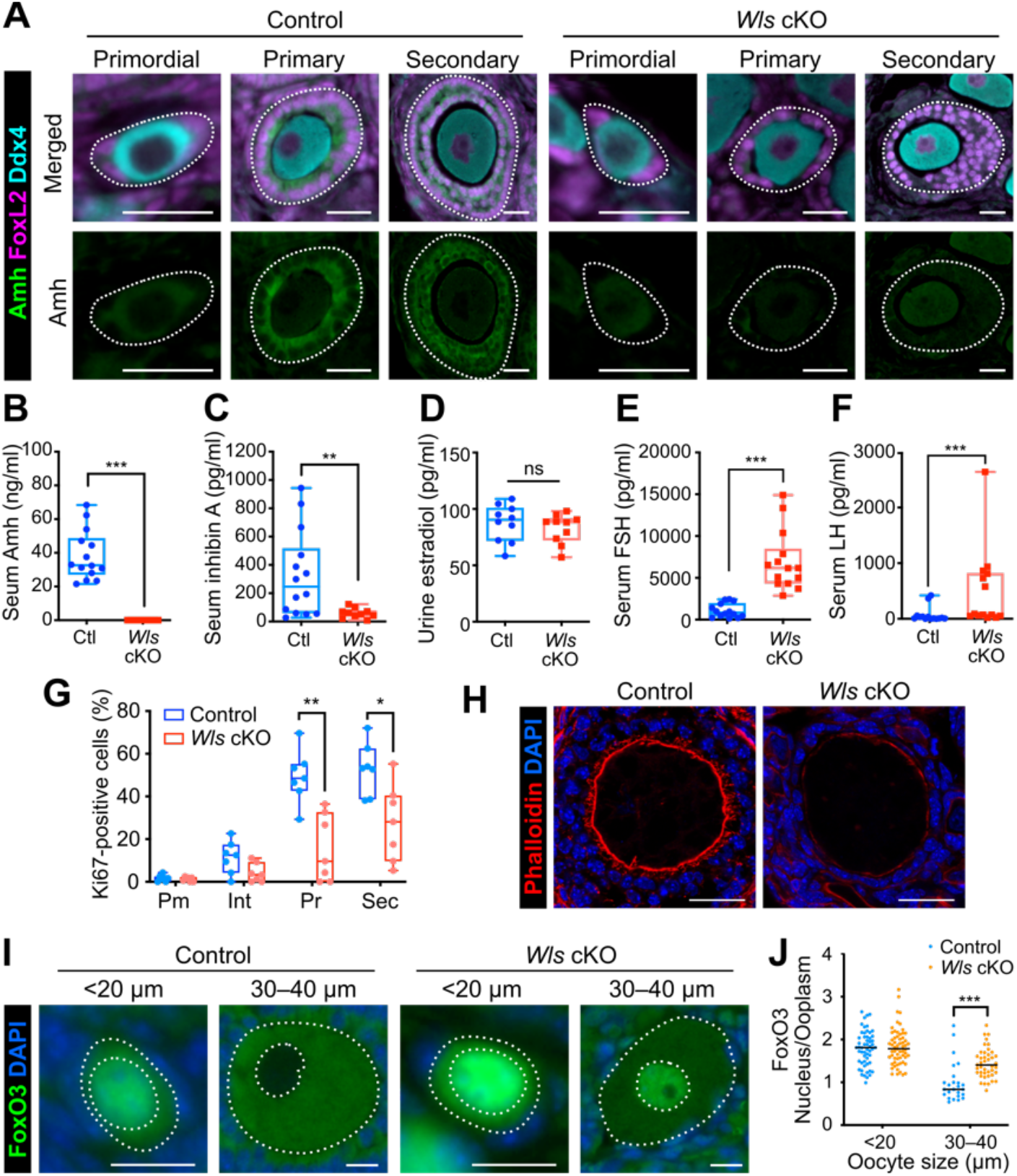
GC differentiation and oocyte activation are suppressed in *Wls* cKO mice. (A) Immunostaining of Amh (green), Ddx4 (cyan), and FoxL2 (magenta) in the ovaries of *Wl*s cKO or littermate control mice at 3 weeks of age. Follicles are demarcated with white dotted lines. Scale bars, 20 μm. (B–F) Levels of Amh (B) and inhibin A (C) in serum, of estradiol in urine (D), and of FSH (E) and LH (F) in serum of 8-week-old *Wls* cKO and control (Ctl) mice (*n* = 10 to 14). The boxes indicate the median and 25th and 75th percentiles, and the whiskers represent minimum and maximum values. \*\**P* < 0.01, \*\*\**P* < 0.001; ns, not significant (nonparametric Mann-Whitney matched-pairs test). (G) Percentage of Ki67-positive pre-GCs/GCs for each follicle type (Pm, primordial; Int, intermediate; Pr, primary; Sec, secondary) in *Wls* cKO and control mice (*n* = 7) at 3 weeks of age as determined by immunofluorescence staining. \**P* < 0.05, \*\**P* < 0.01 (unpaired multiple *t* tests with Holm-Sidak correction). (H) Staining of actin fibers with phalloidin (red) and of nuclei with DAPI (blue) for growing follicles from *Wls* cKO or control mice at 3 weeks of age. Scale bars, 25 μm. (I) Immunofluorescence staining of FoxO3 (green) for oocytes of *Wls* cKO or control mice at 4 weeks of age. Nuclei were counterstained with DAPI (blue). The white dotted lines mark the boundaries of each oocyte and its nucleus. Scale bars, 20 μm. (J) The nucleus/cytoplasm ratio of FoxO3 fluorescence intensity in oocytes was determined from images similar to those in (I). Horizontal lines represent the median (*n* = 25 to 60 oocytes from four mice of each genotype). \*\*\**P* < 0.001 (unpaired multiple *t* tests with Holm-Sidak correction).

To assess whether Wnt signaling might trigger PFA, we quantified the number of follicles per ovary and categorized them by follicle type at 2 weeks of age. The number of primordial follicles in *Wls* cKO mice was similar to that in control mice (Figure 2F), suggesting that Wnt ligands are not the stimulus for PFA per se, otherwise the accumulation of primordial follicles in the mutant ovaries would have been expected. The observation that oocytes larger than 20 μm were present in the ovaries of *Wls* cKO mice (Figure 2D and 2E) also suggested that these cells are capable of initiating a growth response to PFA. However, in contrast to control ovaries, the ovaries of 2-week-old *Wls* cKO mice lacked oocytes with a diameter of 45 to 60 μm. Oocytes with a diameter of >45 μm thus constituted 26.8 ± 4.2% (mean ± SEM) of all oocytes in control females but only 0.5 ± 0.5% of those in *Wls* cKO females (*P* = 0.0286, nonparametric Mann-Whitney matched-pairs test) (Figure 2D). The retardation or arrest of oocyte growth in *Wls* cKO mice therefore appeared to occur between PFA and full maturity. The number of developing follicles was significantly lower in *Wls* cKO mice (Figure 2F), with insufficient GC maturation likely giving rise to follicular atresia.

Wnt signaling plays an important role in female sex determination during embryogenesis (Parma et al., 2006; Vainio et al., 1999). We therefore next examined the effects of postnatal deletion of *Wls* with the use of the *Wt1*^CreERT2^ knock-in allele (Zhou et al., 2008). Control and *Wt1*^CreERT2^;*Wls*^flox/del^ (PN-*Wls* cKO) mice were injected with tamoxifen at P3, P5, and P7 to induce *Wls*^flox^ deletion and were studied at 3 weeks of age (Figure 2G). The phenotype of PN-*Wls* cKO female mice appeared essentially identical to that of *Wls* cKO females. The PN-*Wls* cKO mice thus showed morphologically normal primordial follicles and attenuated GC differentiation in growing follicles (Figure 2H and 2I), indicating that the defect in GC differentiation of *Wls* cKO and PN-*Wls* cKO mice is not the result of disrupted cell fate determination during embryogenesis but rather results from the lack of Wnt signaling during folliculogenesis. The number of primordial follicles in PN-*Wls* cKO mice was also similar to that in control mice (Figure 2J), providing further evidence that initiation of PFA can take place without Wnt signaling. The ovaries of PN-*Wls* cKO mice also showed reduced numbers of growing follicles (Figure 2J), reflecting suppression of folliculogenesis.

### Functional impairment of GC differentiation in *Wls* cKO mice gives rise to insufficient oocyte activation

To evaluate whether GCs in *Wls* cKO mice are functionally mature, we assessed the expression of anti-Müllerian hormone (Amh), a marker for differentiated GCs. Pre-GCs of primordial follicles do not express Amh, with such expression beginning in association with the transition to cuboidal GCs of growing follicles and being followed by the release of Amh into the circulation (Visser et al., 2006). In *Wls* cKO mice, however, immunofluorescence staining revealed only a low level of Amh expression in GCs (Figure 3A). PFA is thought to be a locally regulated process, whereas the later stages of folliculogenesis are influenced markedly by GC-derived paracrine factors and gonadotropins (Sánchez and Smitz, 2012). We therefore next analyzed major GC-derived hormones (Amh, inhibin A, and estradiol) and gonadotropins (follicle-stimulating hormone [FSH] and luteinizing hormone [LH]) in order to investigate GC function and the endocrine system in *Wls* cKO mice. The concentrations of Amh and inhibin A in serum were significantly lower in *Wls* cKO female mice than in controls at 8 weeks of age (Figure 3B and 3C), whereas the urinary concentration of estradiol, which serves to coordinate systemic reproductive functions (Miller and Auchus, 2011; Millier et al., 1994), did not differ between the two genotypes (Figure 3D). These data indicated that GC function is markedly suppressed in *Wls* cKO mice, whereas estradiol production might be controlled independently of Wnt signaling. By contrast, the serum levels of FSH and LH were significantly higher in *Wls* cKO mice than in control mice (Figure 3E and 3F), possibly reflecting a positive feedback response to the suppressed follicle development and lack of ovulation in the mutant females. Pituitary gland function may thus be normal in *Wls* cKO females, even though *Sf1-Cre* is expressed in endocrine glands(Dhillon et al., 2006). Importantly, low Amh and high FSH levels in serum are diagnostic criteria for human POI (Jankowska, 2017; Méduri et al., 2007).

GC proliferation is a key contributor to follicle growth. The mTOR signaling pathway, which is implicated in PFA, also appears to regulate GC proliferation (Yu et al., 2011). To assess GC proliferation in *Wls* cKO mice, we performed immunostaining for Ki67 and measured the signal intensity for all FoxL2-positive GCs within follicles. Whereas the percentage of Ki67-positive GCs increased with follicle growth in both control and *Wls* cKO mice, the increase was less pronounced in the mutant animals (Figure 3G). Most pre-GCs of primordial follicles were negative for Ki67 in both control and *Wls* cKO mice (Figure 3G). Transzonal projections (TZPs) are membranous extensions from GCs that pass through the zona pellucida to the oocyte cell membrane and are important for normal oocyte development (Albertini et al., 2001; Carabatsos et al., 1998). Staining of filamentous actin with phalloidin revealed the absence of obvious TZP structures in *Wls* cKO ovaries (Figure 3H). These results thus indicated that the abrogated folliculogenesis of *Wls* cKO mice is attributable to impaired GC proliferation and the inability of GCs to support oocyte growth.

Although we found that oocyte growth is initiated in *Wls* cKO mice, it was unclear whether the oocytes undergo normal differentiation and activation. To evaluate oocyte status, we analyzed the expression of FoxO3, a transcription factor that contributes to maintenance of oocyte dormancy, by quantifying the nuclear to cytoplasmic ratio of its immunofluorescence intensity (Castrillon et al., 2003). In control mice, whereas primordial follicles manifested a nuclear FoxO3 localization, FoxO3 was exported from the nucleus during PFA (Figure 3I and 3J). However, in *Wls* cKO mice, both oocytes with a diameter of <20 μm and those with a diameter of 30 to 40 μm showed a higher FoxO3 intensity in the nucleus than in the cytoplasm (Figure 3I and 3J). These data thus indicated that, even if oocytes increase in size, they do not undergo the normal activation process in *Wls* cKO mice. They further suggested that activated GCs are necessary to break the dormancy of oocytes.

### The pre-GC layer is expanded by a dominant stable form of β-catenin

To investigate whether Wnt signaling is sufficient for pre-GC differentiation, we generated *Wt1*^CreERT2^;*Catnb*^lox(ex3)/+^ (Catnb-CA) mice, which express a stable form of β-catenin in the somatic lineage of ovaries (Figure 4A). The stabilized β-catenin binds to T cell factor/lymphoid enhancer factor (TCF/LEF) transcription factors, which activate the expression of target genes for canonical Wnt signaling (Harada et al., 1999). Ovaries of (tamoxifen-treated) Catnb-CA mice were similar in size to those of control mice at 3 weeks of age, but they more spherical and had a smoother surface compared with control ovaries (Figure 4B). The observation that somatic cells were densely packed in the ovarian interstitium of Catnb-CA mice suggested that hyperproliferation of interstitial cells was responsible for these differences. Morphological abnormalities were not apparent for GCs in growing follicles of the mutant mice, whereas pre-GCs of primordial follicles were not squamous but cuboidal (Figure 4B). An increase in pre-GC layer thickness was also detected in primordial follicles containing oocytes with a diameter of <20 μm in Catnb-CA mice (Figure 4C), and Ki67 immunostaining revealed that the proliferation of pre-GCs in primordial follicles was enhanced (Figure 4D). Follicles with oocytes of <20 μm and with four or fewer pre-GCs/GCs that showed no obvious columnar shape were classified as primordial follicles in the mutant ovaries. These results thus revealed that Wnt signaling indeed promotes GC differentiation. Of note, Catnb-CA mice showed a normal subcellular localization pattern for FoxO3 in their oocytes (Figure 4E).

**Figure 4.**
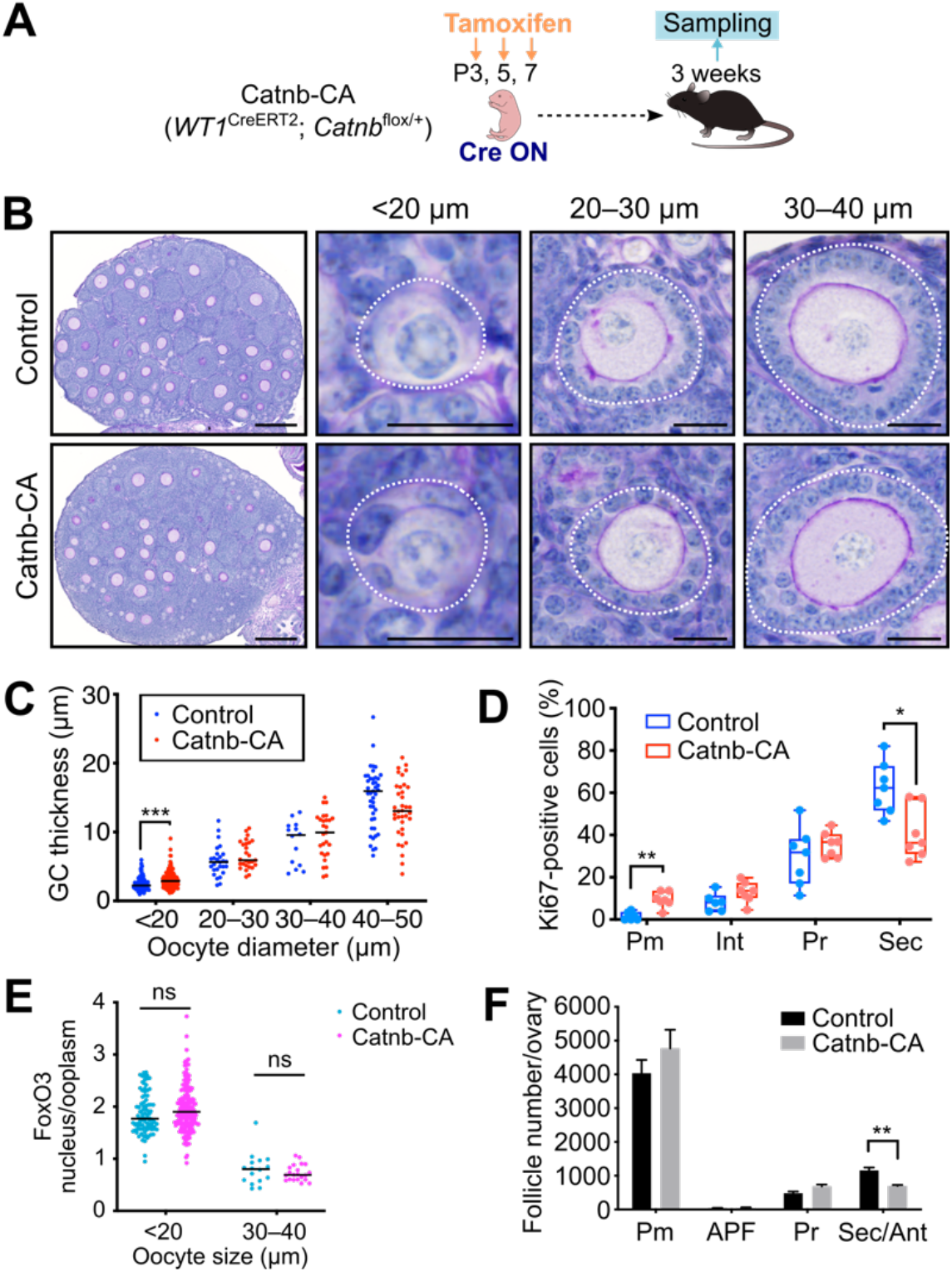
Constitutively active β-catenin promotes pre-GC differentiation and proliferation. (A) Experimental scheme for induction of a constitutively active form of β-catenin in ovarian somatic cells of Catnb-CA mice. (B) PAS-H staining of ovarian sections from (tamoxifen-treated) Catnb-CA and littermate control mice at 3 weeks of age. Follicles were classified by oocyte size and are demarcated by white dotted lines. Scale bars, 200 μm (leftmost panels) or 20 μm (other panels). (C) GC layer thickness categorized by oocyte diameter for Catnb-CA and control mice at 3 weeks of age. Horizontal lines represent the median (*n* = 14 to 140 follicles from five mice of each genotype). \*\*\**P* < 0.001 (unpaired multiple *t* tests with Holm-Sidak correction). (D) Percentage of Ki67-positive pre-GCs/GCs for each follicle type (Int, intermediate) of Catnb-CA and control mice at 3 weeks of age as determined by immunofluorescence staining. Boxes indicate the median and 25th and 75th percentiles, and whiskers represent minimum and maximum values (*n* = 7 mice per genotype). \**P* < 0.05, \*\**P* < 0.01 (unpaired multiple *t* tests with Holm-Sidak correction). (E) The nucleus/cytoplasm ratio of FoxO3 immunofluorescence intensity for oocytes of Catnb-CA and control mice at 3 weeks of age. Horizontal lines represent the median (*n* = 16 to 195 from five mice of each genotype). ns, not significant (unpaired multiple *t* tests with Holm-Sidak correction). (F) Quantification of follicle number per ovary for 3-week-old Catnb-CA and control mice. Follicles were classified as primordial (Pm), activated primordial (APF), primary (Pr), or secondary/antral (Sec/Ant). Data are means + SEM (*n* = 7 mice per genotype). \*\**P* < 0.01 (unpaired multiple *t* tests with Holm-Sidak correction).

Quantification of follicle number revealed no depletion of primordial follicles or increase in the number of developing follicles in Catnb-CA mice (Figure 4F), suggesting that β-catenin stabilization (activation of canonical Wnt signaling) is not sufficient for induction of PFA. Inhibition of secondary follicle growth was apparent in the mutant mice, however, with the number of secondary follicles being significantly reduced (Figure 4F), the GCs secondary follicles were less proliferative (Figure 4D). Constitutive activation of Wnt signaling likely affects GCs, interstitial cells, and Theca cells in such a manner that the survival and growth of secondary follicles are impaired. These characteristics are consistent with the reduced proliferative capacity and cancerous changes of GCs previously observed for mice in which β-catenin or the Wnt agonist Rspo1 was forcibly expressed (Boerboom et al., 2005, 2006; De Cian et al., 2017).

### A Wnt activator rescues the phenotype of *Wls* cKO mouse ovaries in vitro

To verify the phenotype of *Wls* cKO mice, we determined the effects of a Wnt inhibitor in ovarian culture. Ovaries isolated from WT mice at P4 were maintained on membrane cell culture inserts for 6 days by the gas-liquid interphase method in the presence of the Wnt inhibitor IWP2, which blocks Porcupine-mediated palmitoylation and consequent secretion of Wnt ligands (Chen et al., 2009), or of dimethyl sulfoxide (DMSO) as a vehicle control (Figure 5A). The ovaries at the end of the culture period thus corresponded to ovaries at P10 in vivo. PAS-H staining revealed that IWP2 markedly suppressed GC layer development, whereas it had only a minimal effect on primordial follicles with an oocyte diameter of <20 μm (Figure 5B and 5C). We then cultured ovaries from *Wls* cKO or control mice with the Wnt activator CHIR99021 in an attempt to rescue the phenotype of the mutant ovaries (Figure 5D). CHIR99021 activates the canonical Wnt signaling pathway by inhibiting glycogen synthase kinase 3 (GSK3) and thereby stabilizing β-catenin(Bennett et al., 2002). CHIR99021 induced a significant thickening of the GC layer at all assessed follicular stages in both control and *Wls* cKO ovaries (Figure 5E and 5F). Of note, the abnormal flattened morphology of GCs in *Wls* cKO ovaries was completely normalized by CHIR99021 treatment (Figure 5E). These data thus indicated that the function of Wnt signaling in folliculogenesis was evident in vitro, and that a Wnt activator was able to promote follicle growth in a commonly adopted culture system.

**Figure 5.**
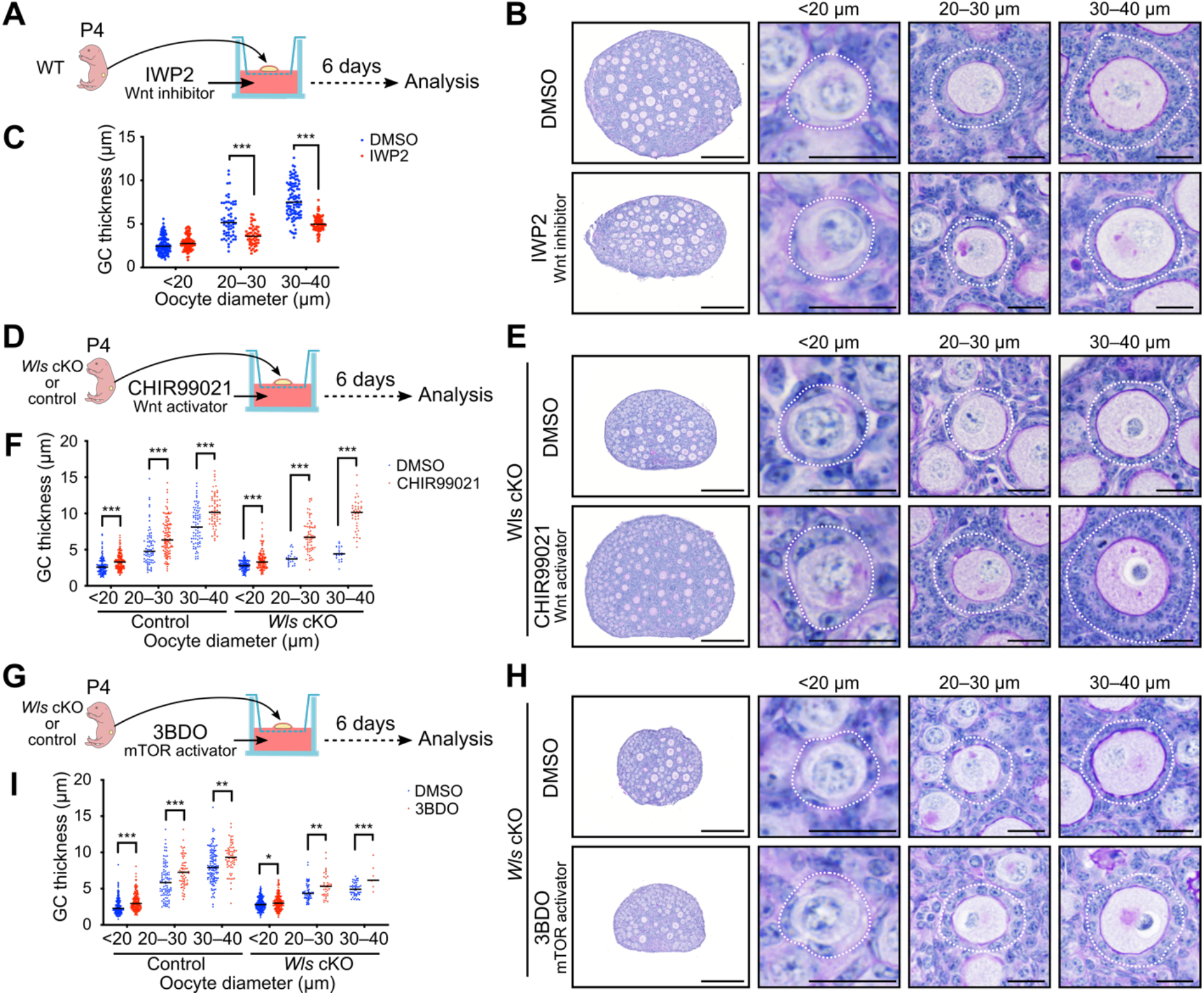
Rescue of the *Wls* cKO ovarian phenotype by a Wnt activator in vitro. (A, D, and G) Experimental design for culture of ovaries from the indicated mice with the Wnt inhibitor IWP2 at 2 μM (A), the Wnt activator CHIR99021 at 5 μM (D), or the mTOR signaling activator 3BDO at 100 μM (G) or with the corresponding concentration of DMSO as a vehicle control (B, E, and H) PAS-H staining of sections of ovaries cultured for 6 days with IWP2 (B), CHIR99021 (E), or 3BDO (H). Follicles were classified by oocyte diameter and are demarcated by the white dotted lines. Scale bars, 200 μm (leftmost panels) or 20 μm (other panels). (C, F, and I) GC layer thickness categorized by oocyte diameter for ovaries cultured with IWP2 (C), CHIR99021 (F), or 3BDO (I). Horizontal lines represent the median (*n* = 7 to 202 follicles from four or five mice of each condition). \**P* < 0.05, \*\**P* < 0.01, \*\*\**P* < 0.001 (unpaired multiple *t* tests with Holm-Sidak correction).

The mTOR signaling pathway is implicated in PFA. Nutritional or other factors are thus thought to activate mTOR signaling in pre-GCs and thereby to stimulate the production of KitL required for oocyte activation (Liu et al., 2014). Given that Wnt signaling has been shown to activate mTOR complex 1 (mTORC1) as a result of inhibition of GSK3 (Inoki et al., 2006), we investigated the potential role of Wnt signaling as an upstream regulator of mTOR signaling in GCs. The addition of an activator of mTOR signaling, 3BDO, to ovarian cultures induced a significant increase in GC layer thickness in follicles of *Wls* cKO and control mice (Figure 5G – 5I). However, this rescue effect for *Wls* cKO ovaries was limited, even in growing follicles (Figure 5H and 5Ixs). These data suggested that Wnt and mTOR signaling contribute to PFA in a coordinated manner, rather than through a simple hierarchical relation.

## Discussion

Our data indicate that canonical Wnt signaling is essential for the transition from primordial to primary follicles. We here propose a two-step model for PFA and functional follicle growth: (1) self-activation of Wnt signaling in pre-GCs induces their transition to GCs, and (2) the differentiated GCs then induce oocytes to exit the dormant state (Figure 6). In the absence of Wnt signaling, the entire GC population showed characteristics specific to pre-GCs, including a squamous shape, hypoproliferative state, limited production of Amh, and lack of TZP formation. Wnt-mediated GC differentiation appears to couple PFA with the nuclear-cytoplasmic shuttling of FoxO3 in oocytes to achieve their reawakening. Given that WntVis signals were undetectable in oocytes, our data do not support the possibility that oocytes directly respond to Wnt ligands produced in GCs by activating the canonical Wnt signaling pathway, although further studies will be required to fully exclude this possibility. Our findings thus underscore the importance of GC-oocyte communication for functional follicle growth and fertility, with oocytes being able to complete maturation and attain their full size only with the support of GCs whose differentiation is dependent on Wnt signaling.

**Figure 6.**
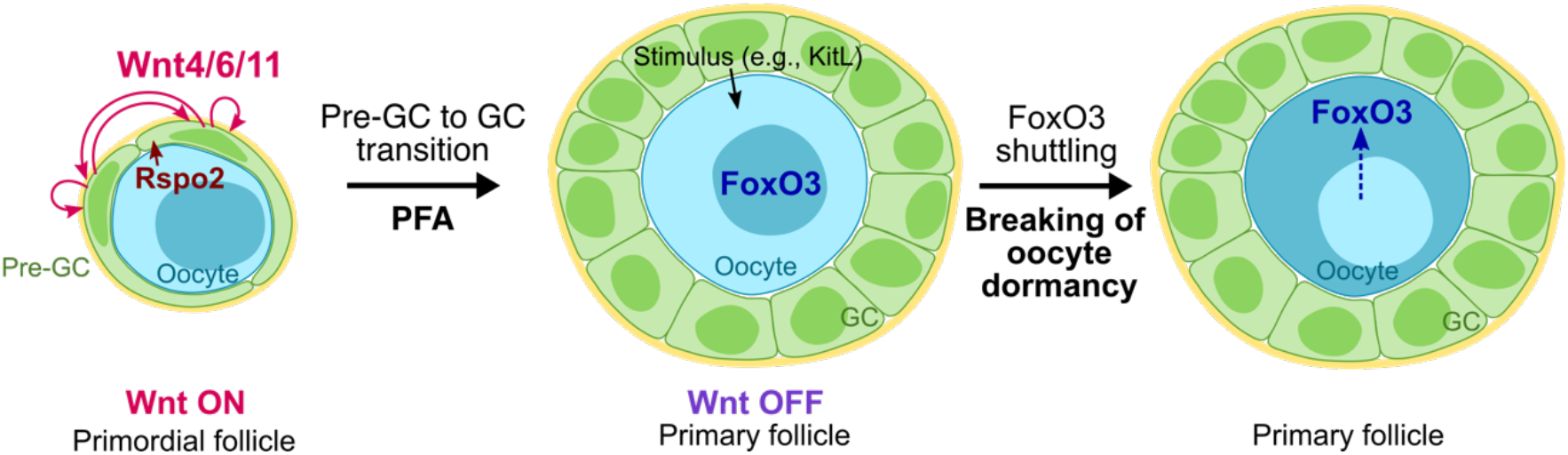
Proposed role of Wnt signaling in folliculogenesis. In primordial follicles, Wnt-responding pre-GCs produce Wnt4, Wnt6, and Wnt11, and oocytes secrete the Wnt agonist Rspo2. The activation of canonical Wnt signaling in pre-GCs promotes their differentiation into GCs during PFA. In primary follicles, differentiated GCs induce the withdrawal of oocytes from their dormant state, as reflected by the translocation of FoxO3 from the nucleus to the cytoplasm.

Our results show that Wnt signal activation occurs exclusively at the primordial follicle stage. Wnt signaling in differentiated GCs is likely detrimental to folliculogenesis, given that forced activation of canonical Wnt signaling in pre-GCs/GCs reduced the number of developing follicles. Our data are consistent with the previous finding that activation of Wnt signaling induced abnormal follicle growth with increased GC apoptosis in an in vitro culture of secondary follicles (Li et al., 2014). The activation of Wnt signaling specifically at the primordial follicle stage is likely achieved as a result of the characteristic expression pattern of *Wnt4/6/11*. Another mechanism may also operate to regulate the timing of Wnt signaling activation, however, given that such activation occurs during a narrower time window compared with that during which these Wnt ligand genes are expressed. It was recently suggested that production of functional Rspo2 by oocytes is important for the activation of canonical Wnt signaling in GCs (De Cian et al., 2020). Given that *Rspo2* mRNA was found to be abundant in oocytes of growing follicles, a mechanism likely exists to inhibit Wnt signaling after the primary follicle stage. BMP15 has been identified as an inhibitor of Wnt signaling during early embryogenesis in *Xenopus* (Di Pasquale and Brivanlou, 2009), and activated mouse oocytes begin to secrete BMP15 at the primary follicle stage (Dube et al., 1998). BMP15 is therefore a candidate mediator of the suppression of Wnt signaling in growing follicles. Wnt signaling in pre-GCs/GCs may thus be precisely controlled at several levels, including the spatiotemporal specificity of Wnt ligand expression and the production of Rspo2 and BMP15 by oocytes.

Recent progress in the field of in vitro gametogenesis (IVG) has had a great impact on reproductive biology and medicine (Hikabe et al., 2016). Fully developed oocytes can now be obtained from embryonic stem cells or induced pluripotent stem cells of mice by the application of IVG techniques. However, the functional integrity of IVG-derived oocytes is lower than that of oocytes in vivo. Somatic cells isolated from embryonic female gonads are required for the growth of oocytes during IVG, but not much attention has been paid to the identity and function of such somatic GCs. In our short-term ovarian culture, a Wnt activator induced GC layer thickening and enhanced oocyte growth. Even though the dormant primordial follicle stage is thought to be skipped in IVG, transient administration of a Wnt activator at the time of follicle formation might be expected to enhance GC differentiation and thereby to increase oocyte integrity. Treatment of ovarian cultures with CHIR99021 increased the thickness of the GC layer in early developing follicles (oocyte size of 20 to 40 μm), whereas such an effect was not observed in response to activation of Wnt signaling in Catnb-CA mice. This difference may be due to a difference in the extent of Wnt signaling activation, or to an effect of CHIR99021 on cell survival (Wang et al., 2015).

In vitro activation (IVA) has recently been described as an innovative method of fertility treatment for women with POI (Kawamura et al., 2013; Suzuki et al., 2015). In this method, the oocyte-awakening process (PTEN-PI3K-Akt-FoxO3 signaling) is targeted in order to activate the few remaining primordial follicles. Wnt-related genes have not been identified as genes responsible for POI, but we have now shown that the hormonal environment of *Wls* cKO mice is similar to that of women with POI (Jankowska, 2017; De Vos et al., 2010). This finding suggests that some cases of POI diagnosed as idiopathic may include those attributable to insufficient GC differentiation. Given that such women are thought to retain intact primordial follicles in their ovaries, the application of Wnt activators to IVA may prove to be clinically beneficial.

## Methods

### Animals

*Sf1-Cre* mice (stock no. 012462), *Wls*^flox^ mice (stock no. 012888), *Wt1*^CreERT2^ mice (stock no. 010912), and *Ddx4-Cre* mice (stock no. 006954) were obtained from The Jackson Laboratory (Carpenter et al., 2010; Dhillon et al., 2006; Gallardo et al., 2007; Zhou et al., 2008). *Wls*^del^ mice, in which the *Wls*^flox^ allele is deleted ubiquitously, were generated by crossing *Wls*^flox^ mice with *Ddx4-Cre* mice. *R26-WntVis* mice (accession no. CDB0303K) were obtained from Laboratory for Animal Resources and Genetic Engineering at the RIKEN Center for Biosystems Dynamics Research, Kobe, Japan (http://www.clst.riken.jp/arg/reporter_mice.html) (Takemoto et al., 2016). *Catnb*^lox(ex3)^ mice were kindly provided by M. M. Taketo (Kyoto University, Japan) (Harada et al., 1999). *Sf1-Cre;Wls*^flox/+^ and *Wls*^flox/del^ mice were used as littermate controls for *Sf1-Cre;Wls*^flox/del^ (*Wls* cKO) mice. Tamoxifen-injected *Wt1*^CreERT2^;*Wls*^flox/+^ and *Wls*^flox/del^ mice were used as littermate controls for *Wt1*^CreERT2^;*Wls*^flox/del^ (PN-*Wls* cKO) mice. Tamoxifen-injected *Wt1*^CreERT2^ and *Catnb*^lox(ex3)/+^ mice were used for littermate control for *Wt1*^CreERT2^;*Catnb*^lox(ex3)/+^ (Catnb-CA) mice. Tamoxifen (0.2 mg per 20 g of body weight) was injected intraperitoneally into mice at P3, P5, and P7. All animal experiments were approved by the Institutional Animal Care and Use Committee of RIKEN. All mouse lines studied were maintained on a mixed genetic background.

### Fertility test

Eight-week-old control or *Wls* cKO female mice (*n* = 7 for each genotype) were housed continuously with WT (C57BL/6N) males for 24 weeks, and the numbers of pups produced were counted.

### In situ hybridization

In situ hybridization was performed with the use of the RNAscope system (Wang et al., 2012). Ovaries from 3-week-old WT mice were fixed in 10% neutral buffered formalin at room temperature for 24 h, dehydrated, and embedded in paraffin. Tissue sections were processed for in situ detection of RNA with the RNAscope 2.5 High Definition (HD)-Red Assay (ACDBio, Hayward, CA). The probes included those for *Dapb* (catalog no. 310043, accession no. EF191515, target region 414–862), *Wnt4* (catalog no. 401101, accession no. NM_009523.2, target region 2147–3150), *Wnt6* (catalog no. 401111, accession no. NM_009526.3, target region 780–2026), and *Wnt11* (catalog no. 405021, accession no. NM_009519.2, target region 818–1643). Sequences of the probes used in Figure S1 are as follows: *DapB* (catalog no. 310043, accession no. EF191515, target region 414–862), *Polr2a* (catalog no. 312471, NM_009089.2, target region 2802–3678), *Wnt1* (catalog no. 401091, accession no. NM_021279.4, target region 1204–2325), *Wnt2* (catalog no. 313601, accession no. NM_023653.5, target region 857–2086), *Wnt2b* (catalog no. 405031, accession no. NM_009520.3, target region 1307–2441), *Wnt3* (catalog no. 312241, accession no. NM_009521.2, target region 134–1577), *Wnt3a* (catalog no. 405041, accession no. NM_009522.2, target region 667–1634), *Wnt5a* (catalog no. 316791, accession no. NM_009524.3, target region 200–1431), *Wnt5b* (catalog no. 405051, accession no. NM_001271757.1, target region 319–1807), *Wnt7a* (catalog no. 401121, accession no. NM_ 009527.3, target region 1811–3013), *Wnt7b* (catalog no. 401131, accession no. NM_009528.3, target region 1597–2839), *Wnt8a* (catalog no. 405061, accession no. NM_009290.2, target region 180–1458), *Wnt8b* (catalog no. 405071, accession no. NM_011720.3, target region 2279– 3217), *Wnt9a* (catalog no. 405081, accession no. NM_139298.2, target region 1546–2495), *Wnt9b* (catalog no. 405091, accession no. NM_011719.4, target region 706–1637), *Wnt10a* (catalog no. 401061, accession no. NM_009518.2, target region 479–1948), *Wnt10b* (catalog no. 401071, accession no. NM_011718.2, target region 989–2133), and *Wnt16* (catalog no. 401081, accession no. NM_053116.4, target region 453–1635).

### Immunostaining and histology

For immunofluorescence staining on paraffin sections, ovaries were fixed overnight at 4°C with 4% paraformaldehyde in phosphate-buffered saline (PBS). The fixed tissue was dehydrated, embedded in paraffin, and then sectioned at a thickness of 5 μm. The sections were depleted of paraffin and rehydrated according to standard protocols. For antigen retrieval, they were incubated either at 110°C for 15 min with citrate buffer (pH 6.0) or at 90°C for 20 min with HistoVT One (Nacalai Tesque, Kyoto, Japan). After washing with PBS containing 0.1% Tween 20 (PBST), the sections were incubated for 1 h at room temperature in a blocking buffer, stained overnight at 4°C with primary antibodies, and then exposed for 2 h at room temperature to a 1:500 dilution of secondary antibodies labeled with Alexa Fluor 488, 568, or 647 (A11057, A21447, A21463, and A10042, Thermo Fisher Scientific; or 703-545-155, Jackson ImmunoResearch). The primary antibodies included chicken anti-GFP (1:500 dilution; GFP-1010, Aves Labs), rabbit anti-Ddx4 (1:500; ab13840; Abcam), mouse anti-Ddx4 (1:200; ab27591, Abcam), goat anti-FoxL2 (1:200; ab5096, Abcam), rabbit anti-Amh (1:100; GTX129593, GeneTex), rabbit anti-FoxO3 (1:500; 12829, Cell Signaling), and rat anti-Ki67 (1:400; 14-5698-82, eBioscience). For immunofluorescence staining on frozen sections, the fixed ovaries were immersed in sucrose gradients (10%, 20%, and 30%) in PBS sequentially at 4 °C, then tissues were embedded in optimal cutting temperature compound (OCT). Frozen samples were sectioned at 6 μm using CryoStar NX70 (Leica Microsystems). For antigen retrieval, cryosections were incubated at 70°C for 30 min with HistoVT One (Nacalai Tesque). After washing with PBS containing 0.1% Tween 20 (PBST), the sections were incubated in blocking buffer for 1 h at room temperature, stained overnight at 4°C with primary antibodies, and then exposed for 2 h at room temperature to a 1:500 dilution of secondary antibodies labeled with Alexa Fluor 488, 568, or Dylight 650 (A10042 and SA5-10029, Thermo Fisher Scientific; or 703-545-155, Jackson ImmunoResearch). The primary antibodies included chicken anti-GFP (1:500 dilution; GFP-1010, Aves Labs), rabbit anti-Cyp17a1 (1:4,000, 14447-1-AP, Proteintech), rat anti-Pecam1 (1:100, sc-18916, Santa Cruz Biotechnology). Antibodies were diluted in blocking buffer or Can Get Signal immunostain solutions (Toyobo, Osaka, Japan). DNA was counterstained with 4′,6-diamidino-2-phenylindole (DAPI). Samples were mounted with Vectashield Vibrance Antifade Mounting Medium (Vector Laboratories). For phalloidin staining, ovaries fixed with 4% paraformaldehyde in PBS were embedded in Tissue-Tek OCT Compound (Sakura Finetek Japan, Tokyo, Japan) and sectioned at a thickness of 6 μm. The sections were washed with PBST, stained with Alexa Fluor 568–conjugated phalloidin (1:100 dilution; A12380, Thermo Fisher Scientific) and DAPI, and mounted.

PAS-H staining was performed according to a standard protocol. In brief, ovaries were fixed in Bouin’s solution, embedded in paraffin, and sectioned at a thickness of 5 μm. The sections were hydrated and treated first with periodic acid solution for 10 min and then with Schiff’s reagent for 15 min. Nuclei were counterstained with hematoxylin.

For quantitative analysis of follicle number, ovaries fixed with Bouin’s solution were embedded in paraffin and serially sectioned at a thickness of 8 μm. After treatment with EDTA buffer (pH 8.0) at 110°C for 15 min, every fifth section was incubated consecutively with antibodies to Ddx4 (1:500 dilution; ab27591, Abcam) and biotinylated secondary antibodies (1:500; BA-1000, Vector Laboratories). Immune complexes were detected with a Streptavidin Biotin Complex Peroxidase Kit (Nacalai Tesque) and Peroxidase Stain DAB Kit (Nacalai Tesque). Nuclei were counterstained with hematoxylin. Only follicles with a visible nucleolus in the oocyte were counted. The raw counts of follicle number were multiplied by 5 to account for the unanalyzed sections and to obtain the estimates of follicle number per ovary. The follicles were classified as primordial follicles (containing an oocyte with a diameter of <20 μm and surrounded by flat pre-GCs), activated primordial follicles (containing an oocyte with a diameter of >20 μm but not containing cuboidal GCs), primary follicles (containing an oocyte surrounded by a single layer of cuboidal GCs), secondary follicles (containing an oocyte surrounded by two or more layers of GCs), and antral follicles (containing an oocyte surrounded by multilayered GCs with antral cavity).

### Image analysis

Immunostaining was examined with a BX53 upright microscope (Olympus) or a slide scanner (Axio Scan.Z1, Zeiss). Phalloidin staining was examined with a confocal laser scanning microscope (TCS SP8, Leica Microsystems). In situ hybridization and PAS-H staining were examined with a slide scanner (Axio Scan.Z1, Zeiss).

For measurement of WntVis or Ki67 signals in pre-GCs/GCs, ovarian sections were subjected to immunofluorescence staining for GFP or Ki67, respectively, as well as for the GC marker FoxL2 and the oocyte marker Ddx4. Nuclei were stained with DAPI. Images were acquired with a BX53 upright microscope (Olympus) or a slide scanner (Axio Scan.Z1, Zeiss). With the use of ImageJ software (NIH), areas positive for both FoxL2 and DAPI were determined as nuclear regions of GCs. The fluorescence intensity of GFP or Ki67 in each region was measured. The lower thresholds for GFP- or Ki67-positive cells were set at the value with 99% accuracy in negative control samples. More than five ovaries for each genotype as well as more than one section per ovary were analyzed. Results were summarized according to follicle type: primordial, intermediate (containing an oocyte surrounded by a mixed single layer of pre-GCs and GCs), primary, and secondary. Only follicles with a visible nucleolus in the oocyte were analyzed.

For quantification of the subcellular localization of FoxO3, ovarian sections were subjected to immunofluorescence staining for FoxO3 and counterstaining with DAPI. Images were acquired with a slide scanner (Axio Scan.Z1, Zeiss) and analyzed with ImageJ software (NIH). The fluorescence intensity of FoxO3 in cytoplasmic and nuclear (DAPI-positive) regions of oocytes was measured together with oocyte diameter. The nuclear to cytoplasmic ratio of FoxO3 intensity was then determined. More than five ovaries for each genotype, and more than one section per ovary, were analyzed.

The thickness of the GC layer was determined as half the difference between the diameters of the follicle and the oocyte as measured in PAS-H–stained ovarian sections with the use of ImageJ software (NIH). More than four ovaries for each genotype or treatment, and more than one section per ovary, were analyzed.

### Measurement of hormone levels

Mice were anesthetized with isoflurane for collection of blood by cardiac puncture, and they were then killed by cervical dislocation. Blood was allowed to clot at room temperature for at least 30 min before centrifugation at 10,000 × *g* for 5 min to obtain serum. For collection of urine, mice were manually restrained and allowed to urinate on disposable plastic trays. Serum and urine were stored at –80°C until analysis. Amh, inhibin A, and estradiol concentrations were measured with enzyme-linked immunosorbent assays (Rat and Mouse AMH ELISA, AL-113, Ansh Labs; Equine/Canine/Rodent Inhibin A ELISA, AL-161, Ansh Labs; and Mouse/Rat Estradiol ELISA, ES180S-100, Calbiotech). Serum levels of FSH and LH were determined with the Luminex method (Oriental Yeast, Tokyo, Japan).

### Ovarian culture

Ovaries from mice at P4 were cultured on Transwell-COL membranes (3.0-μm pore size, Costar) for 6 days by the gas-liquid interphase method (Morohaku, 2019; Morohaku et al., 2016). The basal culture medium comprised α–minimum essential medium supplemented with 10% fetal bovine serum, 1.5 mM 2-*O*-α-D-glucopyranosyl-L-ascorbic acid (Tokyo Chemical Industry), and penicillin (10 U/ml)–streptomycin (10 μg/ml) (Sigma-Aldrich). Ovaries were treated with 2 μM IWP2 (Merck Millipore), 5 μM CHIR99021 (Sigma-Aldrich), or 100 μM 3BDO (Sigma-Aldrich), or with the corresponding concentration of DMSO as a vehicle control. Approximately half of the medium in each well was replaced with fresh medium every other day. The ovaries were maintained at 37°C under 5% CO_2_ and 95% air.

### Statistical analysis

All statistical analysis was performed with GraphPad Prism 8 software. Tests included the nonparametric Mann-Whitney matched-pairs test, unpaired multiple *t* tests with Holm-Sidak correction, and two-way analysis of variance (ANOVA) with Sidak’s post hoc test for multiple comparisons. A *P* value of <0.05 was considered statistically significant.

## Acknowledgments

We thank M. M. Taketo for providing *Catnb*^lox(ex3)^ mice; H. Suzuki for helpful discussion; S. Chunxiao for technical assistance; T. Kato, T. Kimura, K. Minegishi, and K. Miyamichi for comments on the manuscript; and H. Hamada and T. S. Kitajima for critical reading of the manuscript. R.N. is an investigator of the Howard Hughes Medical Institute. This work was supported by the Tomizawa Jun-ichi & Keiko Fund of the Molecular Biology Society of Japan for Young Scientist 2017 as well as by Japan Society for the Promotion of Science KAKENHI grants (17K17690 and 19H05249) to H.M.T.

## Author Contributions

H.M.T. designed the project. O.H., C.Y.L., and H.M.T. performed experiments and analyses. M.K.-A., R.N., and H.M.T. interpreted the data and wrote the paper, with contributions from all other authors.

