## Supplementary material for "Self-activation of Wnt signaling in pre-granulosa cells is required for ovarian folliculogenesis": Figure S1

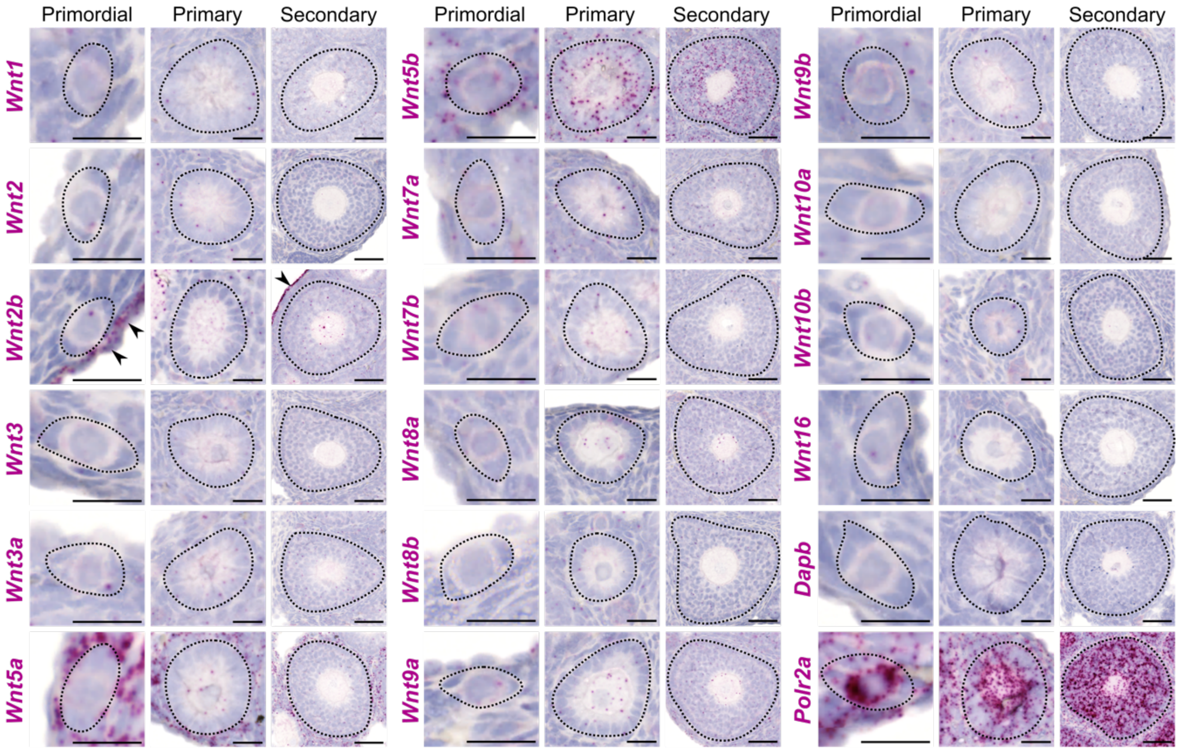


**Figure S1. Expression pattern of Wnt ligands in mouse ovary.** In situ hybridization analysis of *Wnt1*, *Wnt2*, *Wnt2b*, *Wnt3*, *Wnt3a*, *Wnt5a*, *Wnt5b*, *Wnt7a*, *Wnt7b*, *Wnt8a*, *Wnt8b*, *Wnt9a*, *Wnt9b*, *Wnt10a*, *Wnt10b*, *Wnt16*, *Dapb* (negative control), and *Polr2a* (positive control) mRNAs (red) in ovaries of 3-week-old WT mice. Follicles were classified as primordial, primary, or secondary, and they are demarcated with dotted lines. Arrowheads indicate the ovarian epithelium. Scale bars, 50 µm (rightmost panels), or 20 µm (other panels).
