## Supplementary material for "Self-activation of Wnt signaling in pre-granulosa cells is required for ovarian folliculogenesis": Figure S2

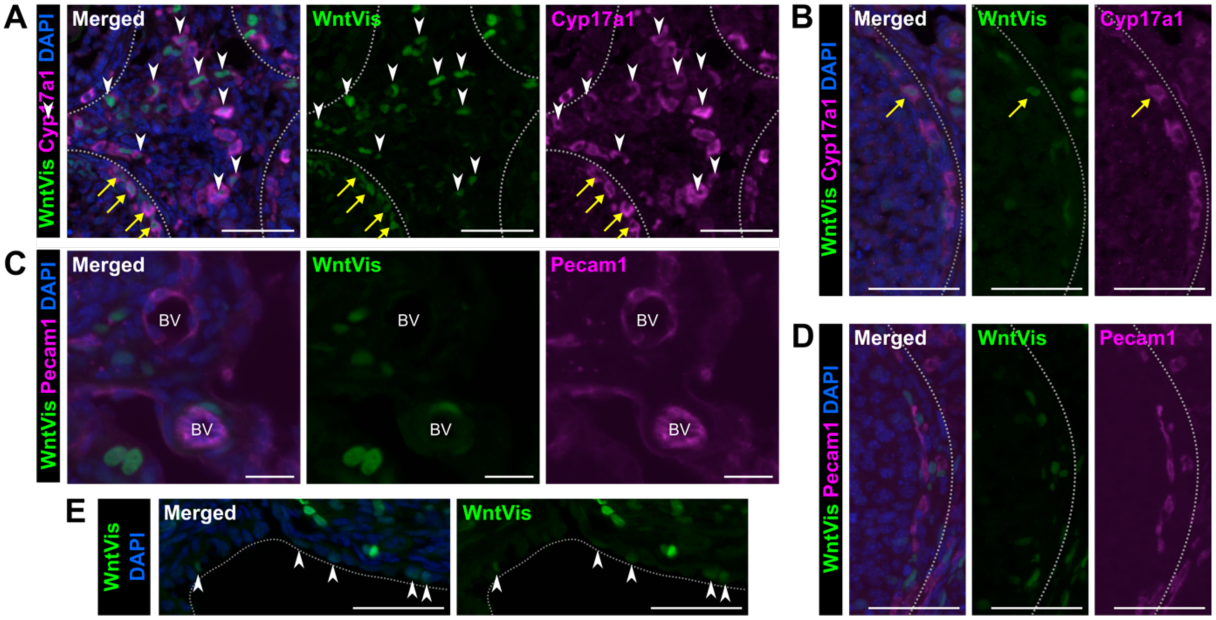


**Figure S2. Interstitial cells, Theca cells, and ovarian epithelium are receiving Wnt signaling occasionally**

(A and B) Immunofluorescence staining of WntVis (green) and Cyp17a1 (magenta) in stromal (A) or Theca cell (B) regions of ovaries from 4-week-old *R26-WntVis* mice. Nuclei were counterstained with DAPI (blue). White arrowheads indicate interstitial cells which are double-positive for WntVis and Cyp17a1. Yellow arrows indicate Theca cells which are double-positive for WntVis and Cyp17a1. The gray dotted lines mark the boundaries of antral follicles. Scale bars, 50 µm.

(C and D), Immunofluorescence staining of WntVis (green) and Epcam (magenta) in blood vessels (C) or capillary vessels around follicles (D) of ovaries from 4-week-old *R26-WntVis* mice. Nuclei were counterstained with DAPI (blue). The gray dotted lines mark the boundaries of antral follicles. BV, blood vessels. Scale bars, 20 µm (C), or 50 µm (D).

(E) Immunofluorescence staining of WntVis (green) in ovarian epithelium of 4-week-old *R26-WntVis* mice. Nuclei were counterstained with DAPI (blue). The gray dotted lines mark the boundaries of ovaries. Scale bars, 50 µm. Arrowheads indicate WntVis-positive ovarian epithelium.
